# Genetic and antigenic variation of foot-and-mouth disease virus during persistent infection in naturally infected cattle and Asian buffalo in India

**DOI:** 10.1101/586644

**Authors:** Jitendra K. Biswa, Rajeev Ranjan, Saravanan Subramaniam, Jajati K. Mohapatra, Sanjay Patidar, Mukesh K. Sharma, Miranda R. Bertram, Barbara Brito, Luis L. Rodriguez, Bramhadev Pattnaik, Jonathan Arzt

## Abstract

The role of foot-and-mouth disease virus (FMDV) persistently infected ruminants in initiating new outbreaks remains controversial, and the perceived threat posed by such animals hinders international trade in FMD-endemic countries. In this study we report longitudinal analyses of genetic and antigenic variations of FMDV serotype O/ME-SA/Ind2001d sublineage during naturally occurring, persistent infection in cattle and buffalo at an organised dairy farm in India. The proportion of animals from which FMDV RNA was recovered was not significantly different between convalescent (post-clinical) and sub-clinically infected animals or between cattle and buffalo across the sampling period. However, infectious virus was isolated from a higher proportion of buffalo samples and for a longer duration compared to cattle. Analysis of the P1 sequences from recovered viruses indicated fixation of mutations at the rate of 1.816 × 10^-2^ substitution/site/year (s/s/y) (95% CI 1.362-2.31 × 10^-2^ s/s/y). However, the majority of point mutations were transitional substitutions. Within individual animals, the mean dN/dS (ω) value for the P1 region varied from 0.076 to 0.357, suggesting the selection pressure acting on viral genomes differed substantially across individual animals. Statistical parsimony analysis indicated that all of the virus isolates from carrier animals originated from the outbreak virus. The antigenic relationship value as determined by 2D-VNT assay revealed fluctuation of antigenic variants within and between carrier animals during the carrier state which suggested that some carrier viruses had diverged substantially from the protection provided by the vaccine strain. This study contributes to understanding the extent of within-host and within-herd evolution that occurs during the carrier state of FMDV.

## Introduction

Foot-and-mouth disease (FMD) is a highly contagious vesicular, viral disease of domesticated and wild even-toed ungulates. The classical clinical FMD syndrome in ruminants is characterised by fever, anorexia, lameness, and vesicles in and around the mouth, feet, and teats. Morbidity can reach 100%, whereas high mortality occurs occasionally amongst young-stock (1-3). The causative agent, FMD virus (FMDV), is the prototype member of the genus *Aphthovirus* in the *Picornaviridae* family (4). The FMDV viral particle contains a single-stranded positive sense RNA genome of approximately 8.2 kb nucleotides in length, enclosed in an icosahedral non-enveloped capsid consisting of 60-copies of each of the four structural proteins VP1, VP2, VP3 and VP4 (5). Seven genetically and antigenically distinct serotypes of FMDV exist (O, A, Asia-1, C, SAT1-3), and within each serotype there are a substantial number of topotypes/genotypes and lineages which have varying degrees of genetic and antigenic diversity (6).

FMDV-infected ruminants typically clear generalized infection within 10 days. However, approximately 50% of FMD-recovered ruminants become FMDV-carriers, defined as animals from which FMDV can be detected in oro-pharyngeal fluid (OPF) more than 28 days post-infection (7-9). The mechanisms that mediate FMDV persistence in specialized regions of nasopharyngeal mucosa are incompletely elucidated, but have been shown to result from a dynamic host-virus interaction at the site of persistence (10-12). Additionally, vaccination does not protect against persistent infection (10, 11, 13), and vaccinated animals often experience neoteric, subclinical infections (14).The duration of FMDV persistent infection may be influenced by a combination of undetermined host and viral factors, and may vary from months to years depending upon the epidemiological context (15-17). The role of persistently infected animals in the evolution and ecology of FMDV remains controversial (7, 18). Although circumstantial evidence from field studies has linked carrier cattle to subsequent outbreaks (19-23), transmission from persistently infected cattle to susceptible naïve animals has not been demonstrated under experimental conditions (24, 25). Yet, oropharyngeal fluid harvested from carriers has been demonstrated to be infectious to naïve cattle (26). Regardless of the epidemiological and physiological basis for risk of transmission from carriers, the perceived risk restricts foreign trade of animals and animal products from endemic regions (27).

Several studies have reported the antigenic and genetic variants of FMDV in the virus population recovered from persistently infected cattle and buffalo under experimental conditions (12, 25, 28-32) or under natural field conditions (14, 17, 21, 33-37). Although within-host genetic variation is common during persistent infection, no consistent genetic changes associated with persistent infection have been identified across studies.

FMDV serotypes O, A, and Asia1 are endemic in India, and serotype O is responsible for more than 80% of FMD outbreaks in the country (2). Under the FMD Progressive Control Program in India, cattle and buffalo (*Bubalus bubalis*) are vaccinated every 6 months; however outbreaks continue to occur throughout much of the country (38). Additionally, many viral strains, including the lineage O/ME-SA/Ind2001d, originated in the Indian sub-continent and have spread to other countries in the Middle East and Southeast Asia (39, 40). Furthermore, some strains of O/ME-SA/Ind2001d isolated in India were found to be antigenically divergent from the vaccine strain, highlighting the importance of vaccine matching and continued monitoring of viruses circulating in the field (41).

The purpose of the current study was to investigate the genetic and antigenic variation of FMDV serotype O/ME-SA/Ind2001d lineage isolated from samples collected sequentially over a period of 13 months from persistently infected cattle and buffalo following a natural FMD outbreak under field conditions on a dairy farm in India. The primary goal was to explore the within-host molecular evolution of persistent FMDV and the role of viral nucleotide variability on the emergence of antigenically variant viruses from persistently infected cattle and buffaloes.

## Materials and methods

### Permissions and ethics

The field outbreak investigations described herein were conducted by federal staff of the Directorate of Foot-and-Mouth Diseases (DFMD) within the Indian Council for Agricultural Research (ICAR), Government of India (GOI) as part of their official duties. All cases described herein occurred spontaneously in domestic livestock with no experimental inoculation or treatment of live animals. No animals were anesthetized or euthanized for the purpose of this study. Sample collection was performed as part of routine field outbreak investigations; samples were subsequently compiled for the sake of the current investigation. As per local standard of operating procedure, ethics approval was not required for the work presented herein. Furthermore no ethics committee exists with oversight of such activities.

### Herd background, FMD outbreak, and case definitions

The herd and epidemiological aspects of the associated 2013-14 FMD outbreak have been described in detail previously (42). Briefly, the current study describes FMDVs derived from samples from an FMD outbreak that occurred at a privately managed, modern dairy farm located in Chattisgarh state, India. The farm was comprised of 4765 cross breed Holstein-Friesian cows, heifers, and Murrah buffaloes. The herd was intensively managed, and the animals were kept in pens which were in close proximity to each other. Animals were routinely vaccinated with a trivalent (Serotypes O, A, and Asia-1) inactivated FMD vaccine four times per year. A presumptive clinical case of FMD was first reported on 24^th^ December 2013, characterised by fever, vesiculo-erosive lesions on the tongue, and inter-digital lesions. Subsequently, the syndrome was definitively diagnosed as FMD by conventional multiplex PCR (mPCR) (43) and antigen-ELISA at the central FMD laboratory, Mukteswar. Additional cases of FMD were recorded for 39 days, and the last case was reported on the farm on 31^st^ January 2014.

All animals were observed daily, and the presence of clinical signs of FMD were determined by the farm’s attending veterinarian. Subsequent to the outbreak, and for the purposes of this study, animals from which FMDV was recovered were classified as either convalescent or subclinical according to the presence or absence of clinical signs of FMD during the outbreak. Convalescent animals had clinical signs of FMD during the outbreak, however all signs of FMD resolved in convalescent animals within 10 days of appearance. Subclinical animals did not have clinical signs of FMD during the outbreak, but were later determined to have been subclinically infected (neoteric subclinical infection) by detection of FMDV or FMDV RNA in OPF. In order to investigate the infection dynamics of FMDV-persistence, OPF samples were collected from 21 convalescent cattle (CC), 16 subclinical cattle (SC), 11 convalescent buffalo (CB), and 6 subclinical buffalo (SB) at 2-3 month intervals for 13-months subsequent to the outbreak. Some of the selected animals were sold during the sampling period, and not all selected animals were available at every time point. The total number of cattle sampled at each time point ranged from 27-29, and the total number of buffalo ranged from 6-15.

### Sample collection and processing

During the acute phase of the outbreak, tissue samples of vesicle epithelium from affected animals were collected and transported to the laboratory in 50% buffered glycerine (pH7.0). These tissue samples were processed as 10% emulsion of homogenised suspension in PBS, and the lysates were centrifuged at 3000g for 15 minutes. The supernatants were used for virus isolation (VI), antigen-ELISA, and extraction of viral RNA for genome amplification. OPF was collected using a probang cup (44) and samples were treated with trichlorotrifluoroethane (TTE) to dissociate the FMDV-antibody complex as previously described (45). All processed OPF samples and clinical sample supernatants (approximately 300µl) were inoculated onto LFBK-αVβ6 cell monolayers for virus isolation, and the remainder of the samples were stored at −70°C for further use.

### FMDV RNA detection

Approximately 500µl of supernatant of clinical sample suspension or OPF prior to TTE treatment was used for the extraction of total RNA using an RNeasy Mini Kit (QIAGEN, Germany). The extracted RNA was quantified using Nanodrop spectrophotometer (ThermoScientific, USA) and reverse transcribed using MMLV reverse transcriptase enzyme (Promega, USA) and oligo d(T)_15_ primer. To improve the sensitivity and specificity of FMDV RNA detection, samples were analysed using both serotype-differentiating agarose gel electrophoresis-based mPCR and SYBR green rRT-PCR, with results interpreted in parallel (samples were considered positive if they were positive on either test). The assays were performed as previously described (43).

### Virus isolation, genome amplification and sequencing

Virus isolation was carried out using the LFBK-αVβ6 cell line (46) through serial cell passage. Supernatant from samples with cytopathic effect in LFBK-αVβ6 cells was clarified and stored at −80°C for subsequent use. For genome amplification and sequencing, the total RNA was extracted from the low-passage infected cell culture supernatant (500 µl) using an RNeasy Mini Kit (QIAGEN, Germany), and cDNA synthesis was carried out using an oligo d(T)_15_ primer and MMLV reverse transcriptase (Promega, USA) enzyme. The structural protein coding region (P1) was amplified and sequenced on an ABI 3130 DNA analyser (Applied Biosystems, USA) as previously described (41). Multiple sequence reads were assembled using EditSeq module (Lasergene 10, DNAStar Inc., USA) and analysed using MEGA 6.06 (47)software. Sequences recovered in this study were submitted to GenBank (accession #MG893512 –MG893552).

### Sequence analyses

The P1 sequences obtained in this study and related FMDV O/ME-SA strains obtained from GenBank were aligned using Clustal W. A maximum likelihood phylogenetic tree was constructed using GTR nucleotide substitution model and 10,000 bootstrap replicates implemented in MEGA 7. The genetic distance between sequences was computed using the p-distance implemented in MEGA7. For identification of codons under selection pressure, sequences of both carrier and outbreak viruses obtained in this study were analysed by Single Likelihood Ancestral Counting (SLAC), Fixed-Effects Likelihood (FEL), and Internal FEL (IFEL) methods using the best fit nucleotide model estimated with HyPhy (48). The sites under episodic diversifying selection were detected using Mixed Effect Model of Evolution (MEME) (48). In order to trace virus movement, statistical parsimony analysis was carried out using network estimation implemented in TCS v1.21software (49) with a cut-off of 90%. To estimate the nucleotide substitution rate, the phylogeny was constructed using Bayesian methods implemented in BEAST 1.8.4 (50). The evolutionary rate was calculated using the relaxed uncorrelated lognormal clock and exponential population size model under Bayesian Markov chain Monte Carlo method implemented in BEAST 1.8.4 (50).

### Antigenic analyses

In order to determine the antigenic relationship (r-value) between the field virus strains and the vaccine virus strain, a two-dimensional virus neutralization assay (2D-VNT) was performed as previously described (51) using bovine vaccinate serum (BVS) against the currently used FMDV vaccine strain O/IND/R2/1975. Detection of cytopathic effect (CPE) on the LFBK-αVβ6 cell monolayer was used as an indicator system in the neutralization assay. The serum titre was calculated from the regression data as the log_10_ reciprocal serum dilution required for neutralization of 100TCID_50_ of virus (both homologous and heterologous) in 50% of the wells. The one-way antigenic relationship (r-value) was calculated as the ratio between the neutralizing serum titre against the heterologous virus (field strain) to the neutralizing serum titre against the homologous virus (vaccine strain). The test was repeated three times and the mean serum titre was used for the calculation of r-value. The r-values in the range of 0.3-1.0 indicate that the field virus is antigenically homologous to the vaccine strain and therefore the vaccine strain is likely to confer protection against challenge with that specific field virus (52).

### Statistical analysis

The dynamics of persistent infection in cattle in this herd have been reported elsewhere (42). The proportion of carrier animals as determined by PCR and by VI was compared between species and between asymptomatic and clinically affected animals at each time point using the chi-squared test. Additionally, the proportion of carrier animals was compared between consecutive time points using the chi-squared test. Statistical analyses were performed using R (53).

## Results

### Duration of persistent infection

FMDV RNA was detected in OPF samples from cattle and buffalo throughout the study. In both cattle and buffalo, the proportion of carriers was not significantly different between convalescent (post-clinical) and subclinical animals (Fig 1). In cattle, the proportion of FMDV RNA positive animals was 100% at 3 months post-outbreak, but fell significantly (χ^2^=6.58, df=1, p=0.01) to 72% at 5 months post-outbreak, and remained approximately 70% at 7 and 10 months post-outbreak. However, a large proportion of cattle apparently cleared the infection between 10 and 13 months post-outbreak, as FMDV RNA was recovered from significantly fewer (7%;χ^2^=18.8, df=1, p<0.0001) cattle at 13 months post-outbreak (Fig 1A). In buffalo, the proportion of FMDV RNA positive animals was 100% at 3 months post-outbreak, and remained >90% from 5 to 10 months post-outbreak. Similar to the trend in cattle, the proportion of FMDV RNA-positive buffalo fell significantly (χ^2^=5.76, df=1, p=0.02) to 17% at 13 months post-outbreak (Fig 1B). Overall, the proportion of animals from which FMDV RNA was detected tended to be higher in buffalo compared to cattle, however the difference was not significant.

**Fig 1.**
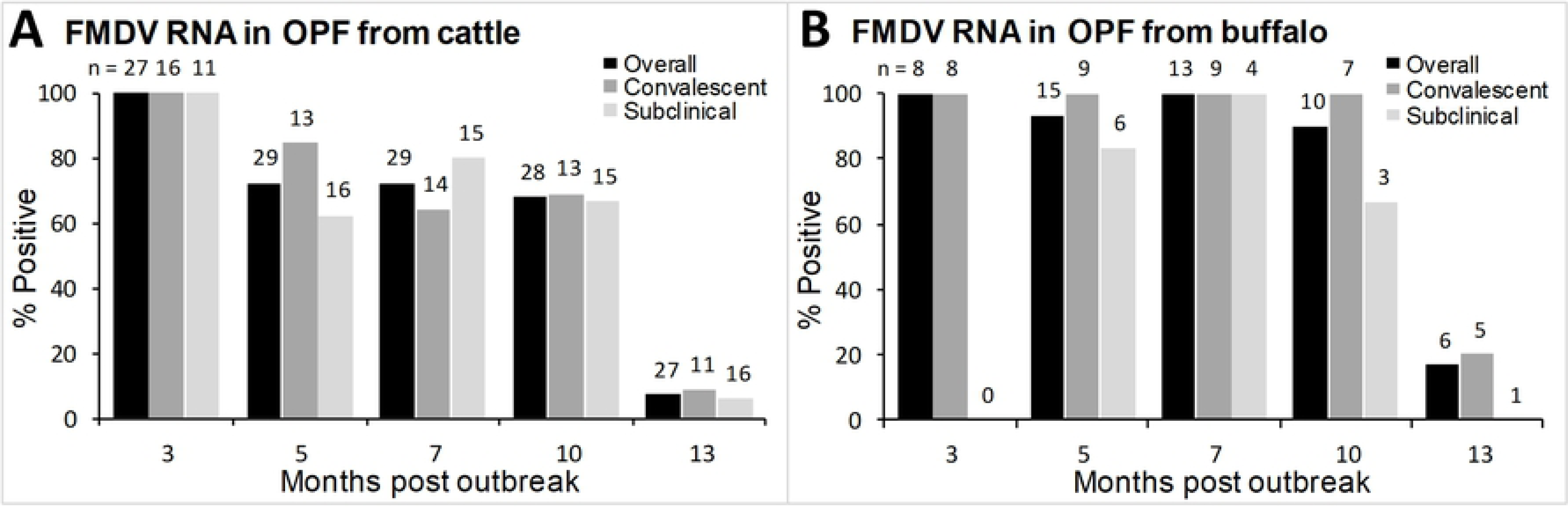
Proportion of convalescent (post-clinical), subclinical, and overall proportion of carrier cattle (A) and buffalo (B) from which FMDV RNA was detected in OPF. Numbers indicate the total number of animals sampled at each sampling period.

Infectious FMDV was isolated from OPF samples from cattle through 7 months post-outbreak and from buffalo throughout the study (Fig 2). Similar to FMDV RNA detection, the proportion of carriers as determined by VI was not significantly different between convalescent and subclinical animals. FMDV was isolated from 59% of cattle at 3 months post-outbreak, however the proportion of VI-positive cattle fell significantly (χ^2^=12.82, df=1, p=0.0003) to 10% at 5 months post-outbreak. FMDV was isolated from 3% of cattle at 7 months post-outbreak, and virus was not isolated from any cattle after 7 months post-outbreak (Fig 2A). In buffalo, FMDV was isolated from 87% of animals at 3 months post-outbreak. Similar to the trend in cattle, the proportion of VI-positive buffalo fell to 40% at 5 months post-outbreak, however the decrease between the 3 & 5 month time points was not significant. The proportion of VI-positive buffalo fell to 16% (*n*=6) at 13 months post-outbreak (Fig 2B). The proportion of VI-positive animals was higher in buffalo compared to cattle, and the difference was significant at 7 and 10 months post-outbreak (χ^2^=8.91, df=1, p=0.003; χ^2^=5.46, df=1, p=0.02, respectively).

**Fig 2.**
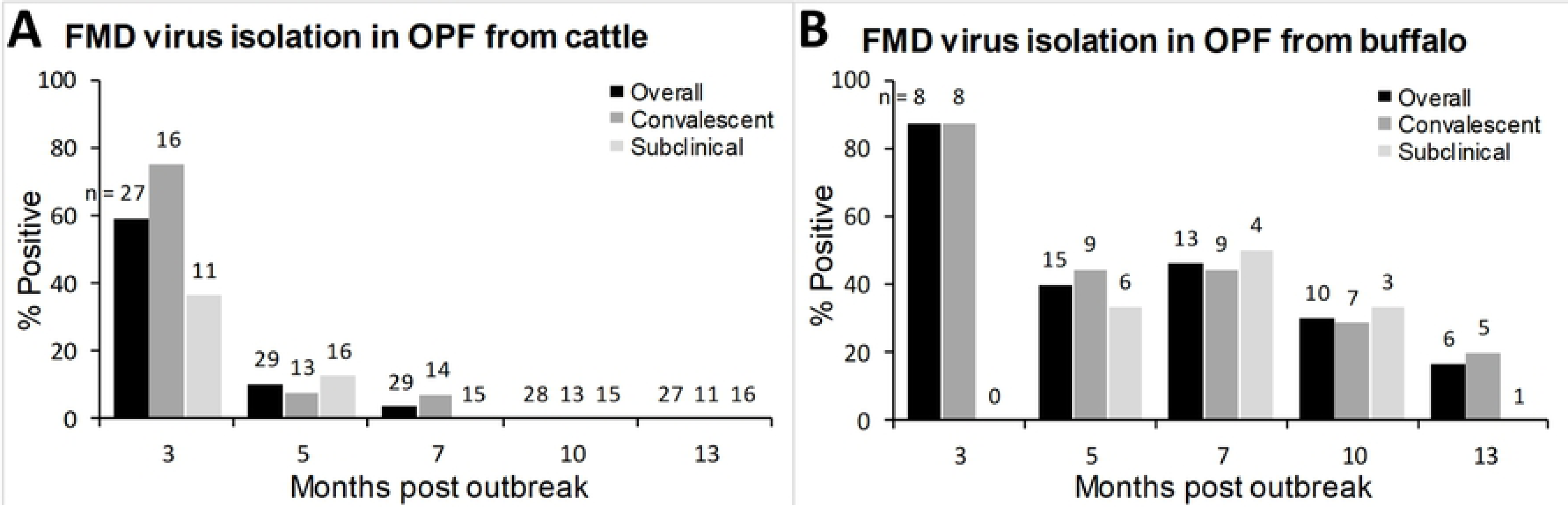
Proportion of convalescent, subclinical, and overall proportion of carrier cattle (A) and buffalo (B) from which infectious FMDV was isolated from OPF. Numbers indicate the total number of animals sampled at each sampling period.

### Phylogenetic analysis and genomic variation of FMDV isolates

FMDV was isolated from a total of 43 OPF samples, of which the FMDV P1capsid-coding region sequence was successfully obtained from 37 samples (86%). An additional four P1sequences were acquired directly from samples of vesicular epithelium collected during the clinical phase of the outbreak during January 2014, for a total of 41 sequences recovered in the current study. The 4 isolates collected during the clinical phase were identical to one another.

The 37 sequences recovered from OPF samples were collected during the carrier phase from 27 different animals. Twenty-seven sequences were recovered from convalescent carrier animals (10 sequences from 9 cattle, 17 sequences from 10 buffalo) and 10 sequences were from subclinical carrier animals (6 sequences from 4 cattle, 4 sequences from 4 buffalo) which had never had clinical signs of FMD. Multiple isolates were recovered from 8 animals.

In the maximum likelihood (ML) phylogenetic tree based on P1-coding region, all sequences recovered in the current study were confirmed to align within sublineage O/ME-SA/Ind2001d (Fig 3). Sequences recovered from carrier buffalo clustered separately from sequences recovered from cattle. Additionally, one cattle-derived sequence (CC20213/AUG/2014) clustered separately from all other isolates, but was more closely related to the other sequences recovered in the current study than to the reference sequences.

**Fig 3.**
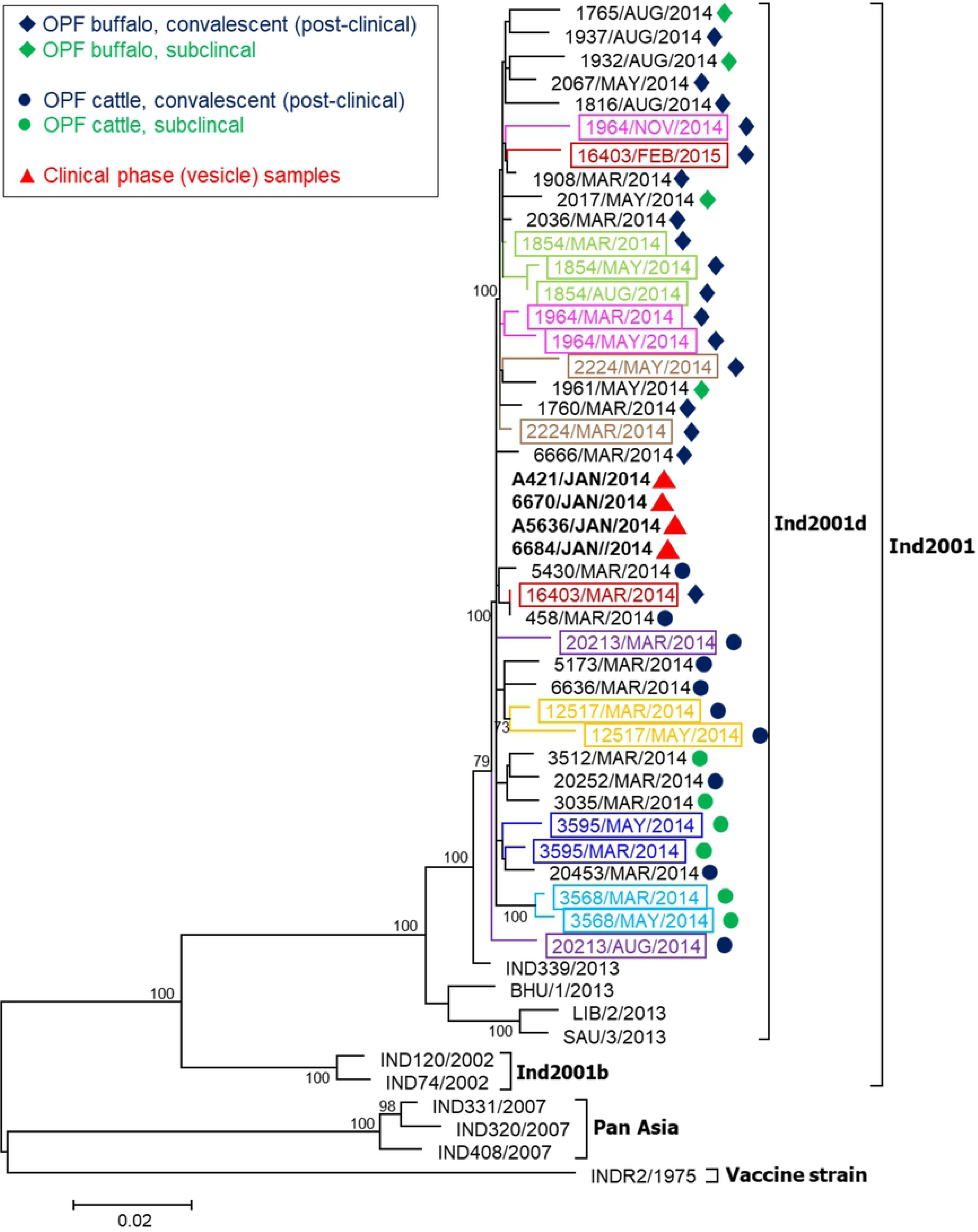
Phylogenetic tree estimated using the maximum likelihood method for the P1-capsid coding region of outbreak (acute) and carrier FMDV isolates. Bootstrap values (>70%, out of 10,000 replicates) are shown near the nodes. Coloured outlines and text denote samples from the same animal.

When comparing viruses from carrier animals to the virus collected during the clinical phase, nucleotide-divergence varied from 0.1% (March 2014, Animal CB1854) to 1.3% (May 2014, Animal CC12517) (Table 1). However, when comparing P1 sequences amongst carrier viruses, a maximum nucleotide (nt) divergence of 2.4% was identified between sequences collected in May 2014 (Animal CC12517) and November 2014 (Animal CB1964), and also between sequences collected in May 2014 (Animal CB2224) and Aug 2014 (Animal SB1932).

**Table 1.**
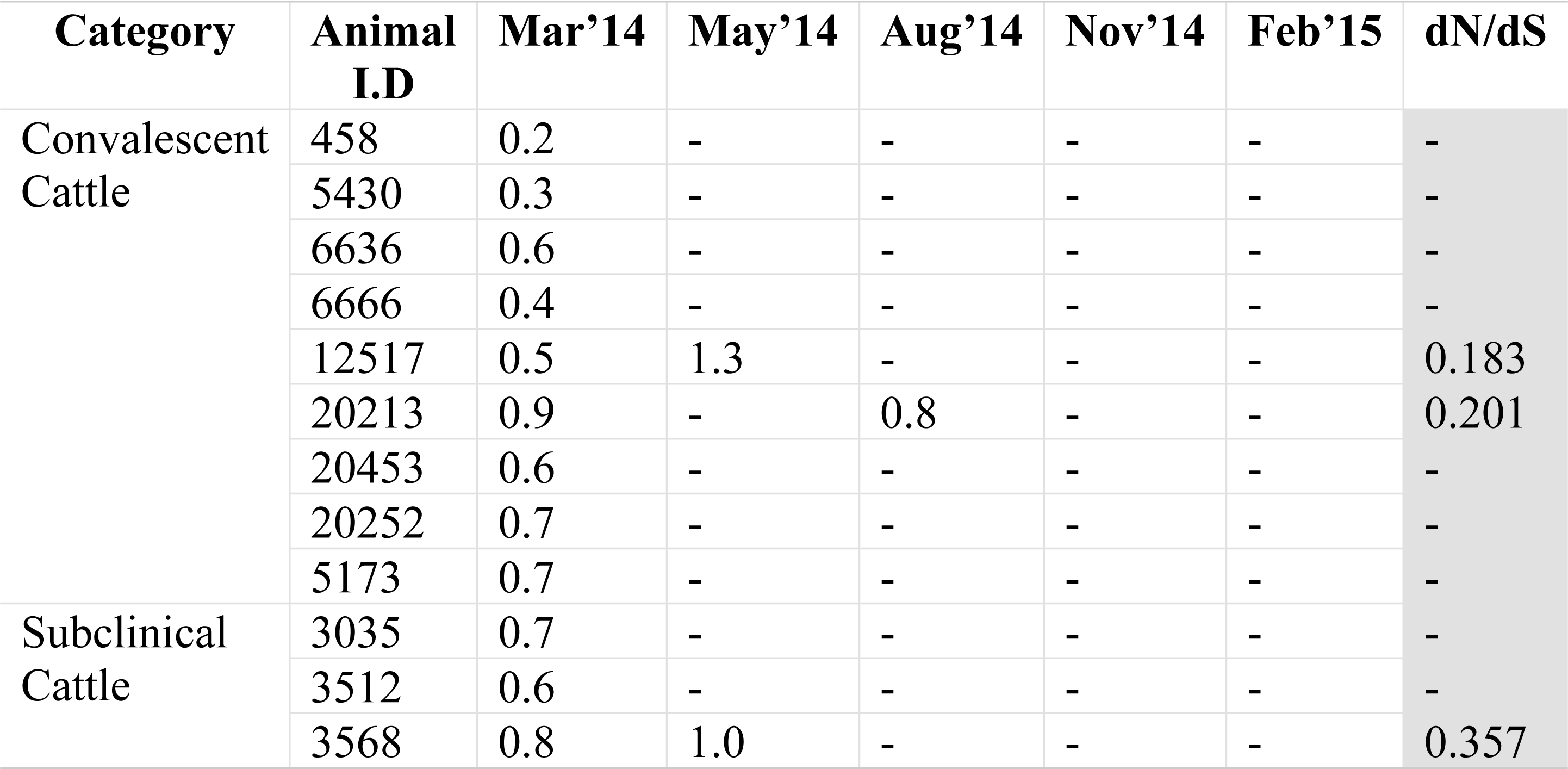
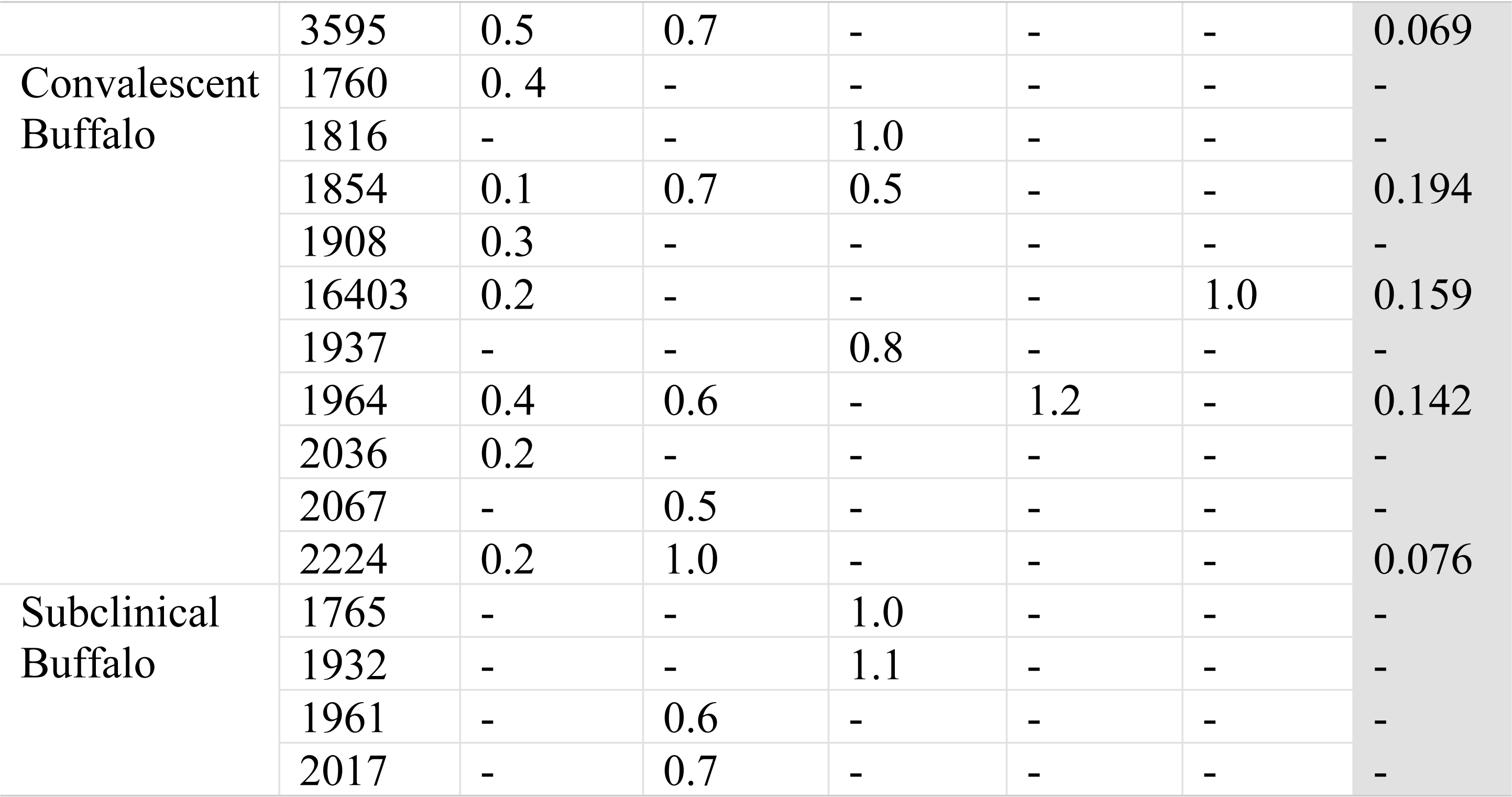
Nucleotide divergence (%) in the capsid coding region of virus isolates from carrier animals compared to the virus isolate collected during the acute phase of the outbreak in January 2014. The dN/dS ratio is reported for animals from which multiple virus isolates were recovered.

Overall, in the capsid coding region, out of 2208 nts, a total 304 sites (13.8%) were found to be polymorphic, of which 208 sites had single nt polymorphism (68.4%) and 96(31.6%) sites had multiple variants. Only point mutations were observed, with no insertion or deletion. As expected, of the total base changes, 86% were transitions and 14% were transversions. The majority (76%) of mutations were synonymous (silent), however 24% of the base substitutions resulted in amino acid (aa) changes in the carrier virus compared to the outbreak virus (Fig 4). The capsid protein coding segments in carrier viruses had variations at 75 (10.2%) aa positions, of which 66 (88%) positions were occupied alternately by two aa and 9 (12%) positions were substituted by more than two aa (Table 2). Out of those 75 positions, 30 sites were located in VP1, 23 sites in VP3, and 17 sites in VP2.

**Table 2.**
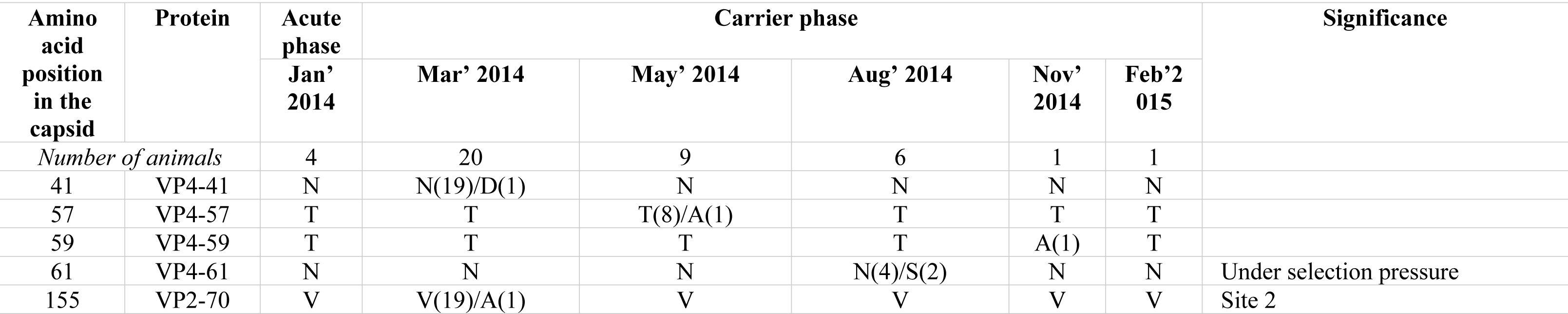
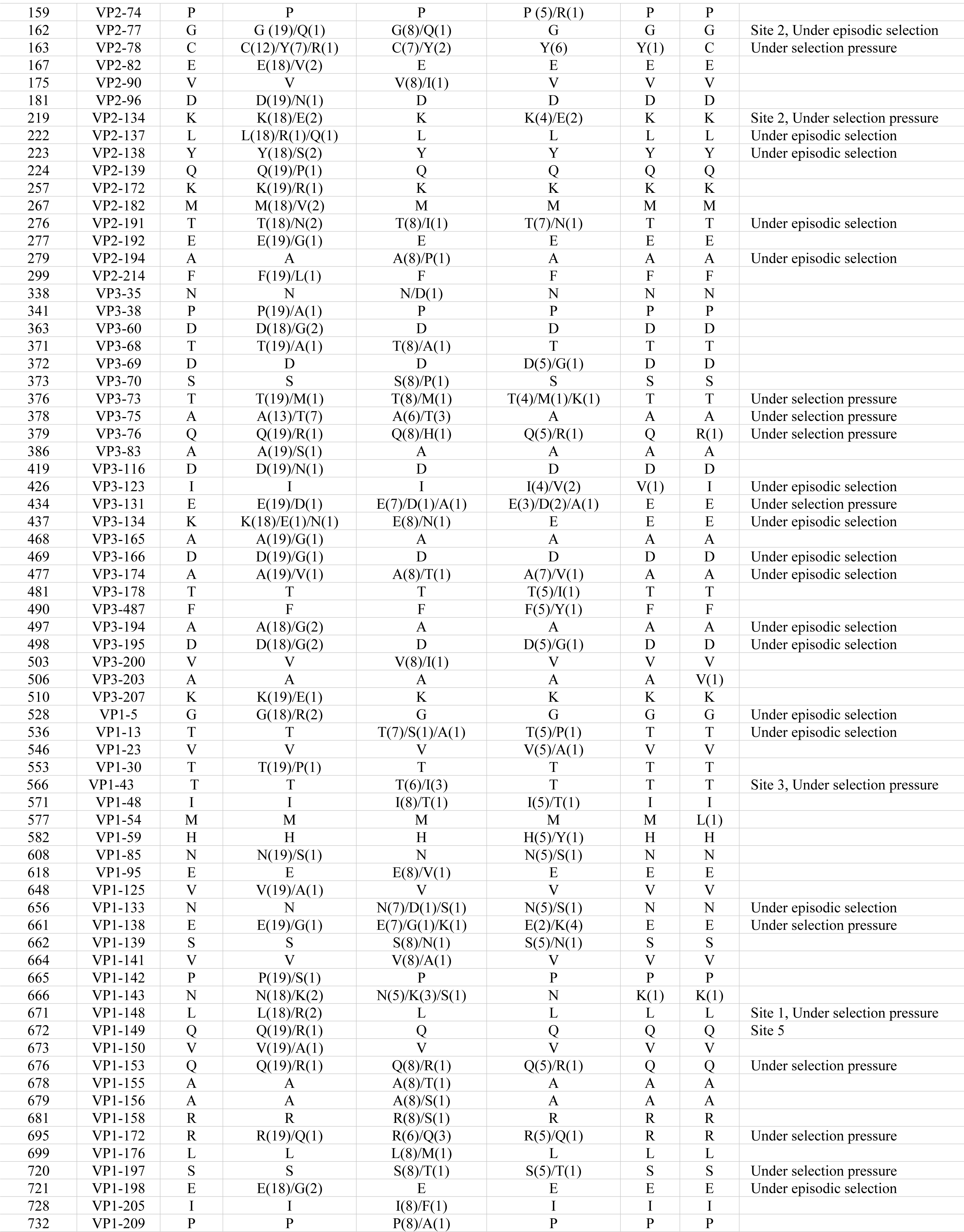
Amino acid variations in the capsid coding region of carrier virus isolates compared to the outbreak virus collected in January 2014. Number of virus isolates in which the changes occurred is in parentheses.

**Fig 4.**
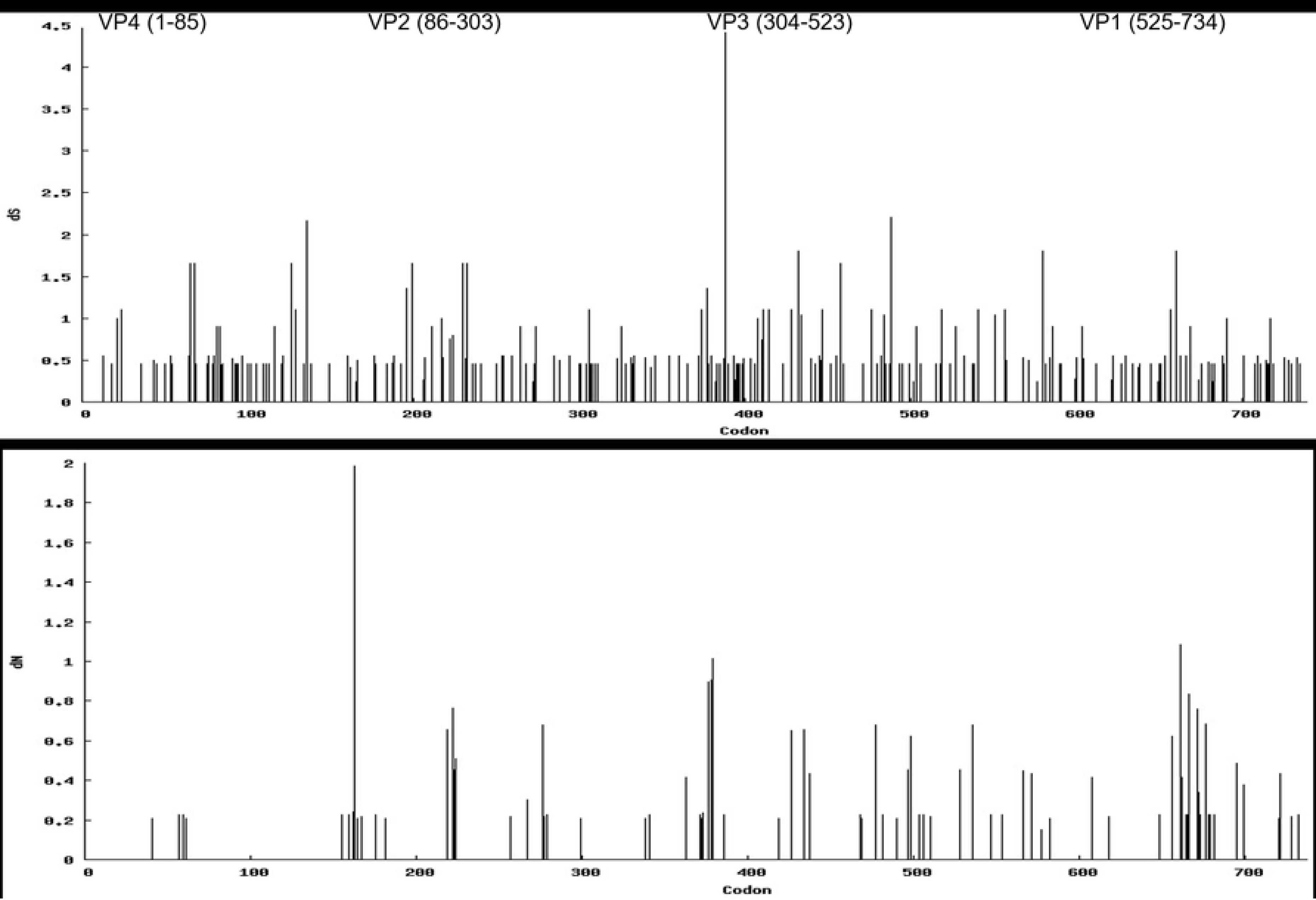
Synonymous (dS; top) and non-synonymous (dN; bottom) changes in the P1-capsid coding region of the genomes of the FMD viruses obtained from carrier animals.

Among the 8 carriers from which multiple sequences were obtained, there were 25 nucleotide substitution sites shared across at least 2 carriers, and a synonymous nucleotide substitution occurred at nucleotide position 1164 (in the VP3 coding region) in 5 carriers (Table 3). These consistent changes across animals may be candidate markers of virus adaptation during their persistent phase.

**Table 3.**
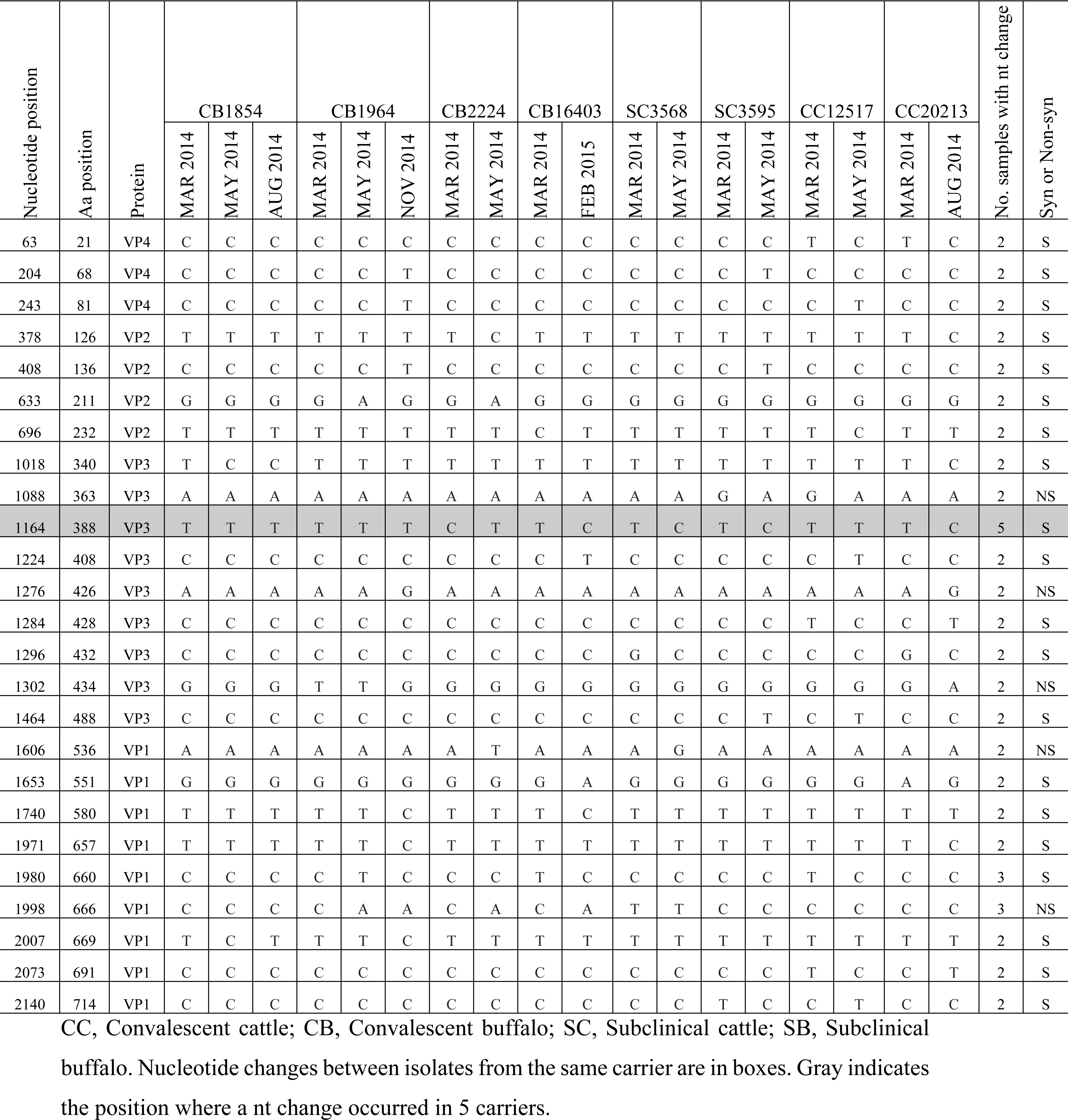
Nucleotide changes that occurred at the same position in at least 2 of the 8 carriers from which multiple sequences were recovered.

### Selection pressure in the capsid coding region of viruses isolated from persistently infected animals

The mean non-synonymous (dN) to synonymous (dS) value (ω) of the entire capsid coding region was found to be 0.188, indicating purifying selection attributable to evolutionary constraints. Across the individual animals for which multiple sequences were available, ω varied from 0.076 to 0.357 (Table 1), suggesting different selective pressure exerted on viral genomes were differed between individuals. The SLAC, FEL, and IFEL methods identified two codons in VP1 (138 and 148) and one codon each in VP2 (78) and VP3 (76) to be under positive selection with statistical significance (Table 4). Only codon 148 was within a known antigenic site, the G-H loop of VP1. Additionally, codon 73 in VP3 was found to be under selection pressure by SLAC and FEL. A total of 28 codons were found to experience episodic diversifying selection, of which 10 codons were in VP3, 9 were in VP1, and 8 were in VP2. Additionally, codon 61 in the highly conserved VP4 region was under episodic selection.

**Table 4.** Amino acid sites identified to be under positive selection in different viral proteins (VP1-VP4) by different site specific models (p < 0.25).

| Method | VP1 | VP2 | VP3 | VP4 |
| --- | --- | --- | --- | --- |
| SLAC | 138 and 148 | 78 | 73 and 76 | - |
| FEL | 138 and 148 | 78 | 73 and 76 | - |
| IFEL | 43, 138, 153, 172 and 197 | 78 and 134 | 68, 75, 76, 131 and 134 | 61 |
| MEME | 5, 13, 43, 133, 138, 148, 153, 172 and 198 | 77, 78, 82, 134, 137, 138, 191 and 194 | 73, 75, 76, 123, 131, 134, 166, 174, 194 and 195 | 61 |

### Statistical parsimony analysis

Multiple phylogenetic analyses were used to improve visualization and inferences of ancestral relationships amongst the viral sequences obtained from individual animals and samples. The root node in the parsimony analysis was formed by four sequences collected during the clinical phase of the outbreak (Fig 5). Isolates recovered from cattle clustered separately from isolates recovered from buffalo. In general, viruses from buffalo had more SNPs relative to the root node compared to viruses from cattle. The nucleotide differences of carriers’ OPF viruses relative to the outbreak virus ranged from a minimum of 3 nt (Animal CB1854, March 2014) to maximum of 28 nt (Animal CC12517, May 2014). Viruses recovered from the same animal at distinct time points generally were more genetically similar than viruses recovered from distinct hosts. However there were exceptions to this trend including a genealogical relationship between viruses isolated from the convalescent carrier buffalo Animal CB1854 in March2014 and two subclinical carrier buffalo in August 2014 (Animal SB1765 and Animal SB1932). An additional noteworthy exception was a convalescent buffalo (Animal CB16403) which had a virus that was similar to the cattle cluster in March 2014; however, the virus isolated from this animal in Feb 2015 clustered with the other viruses derived from buffalo.

**Fig 5.**
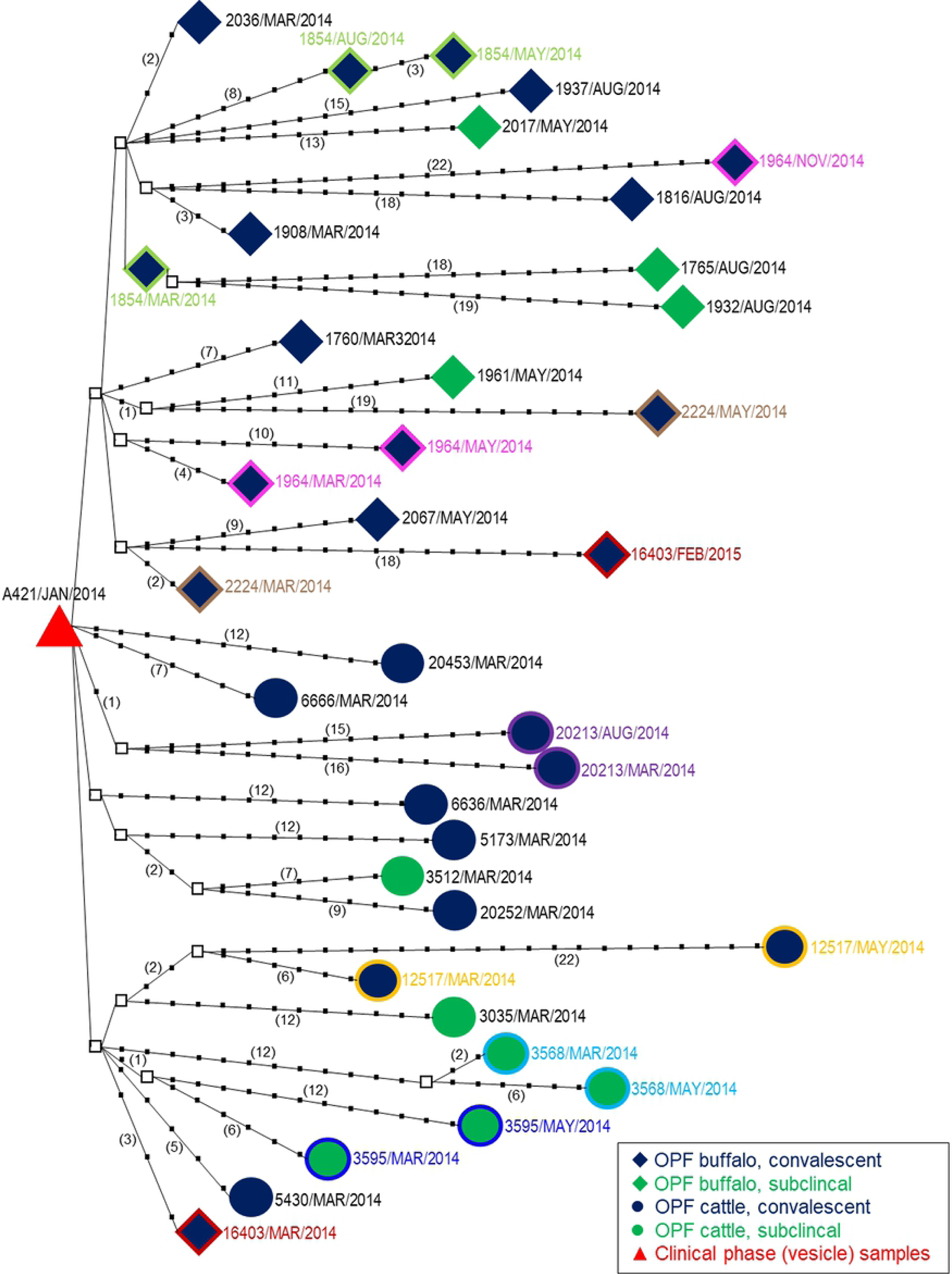
Statistical parsimony analysis of P1-capsid coding regions of outbreak and carrier viruses. The analysis was implemented in TCS v 1.21. Tick marks and numbers in parentheses represent the number of nucleotide changes between the putative ancestors and the virus isolates. Internal nodes (squares) represent un-sampled intermediate sequences inferred by TCS. Coloured outlines and text denote samples from the same animal.

### Evolutionary rate in the P1coding segment of FMDV carrier isolates

Enforced strict and relaxed (Log-normal and Exponential) molecular clocks were used in order to determine the evolutionary rate in the P1 segment of carrier viruses. The Bayes Factor analysis favoured the relaxed log-normal clock. Using the relaxed log-normal clock model, the mean nucleotide substitution was estimated at 1.816 × 10^-2^ substitution/site/year (s/s/y) with a 95% credibility interval of1.362-2.31 × 10^-2^ s/s/y. The coefficient of variation was 0.347, indicating significant rate heterogeneity among branches, and supporting use of the relaxed clock model. The average mutation rate of codon positions 1+2 and 3 was estimated to be 0.519 and 1.963, respectively, indicating a higher contribution of synonymous mutations to the mean evolutionary rate and suggesting the existence of strong constraints for fixation of non-synonymous amino acid mutations due to the need to maintain the functional FMDV capsid structure.

### Antigenic variation in FMDV carrier virus isolates

The antigenic characteristics of outbreak and carrier-derived FMD viruses in relation to the current in-use vaccine strain O/IND/R2/1975 was determined using 2D-VNT (Table 5). The outbreak virus (C6670 and C5636) had an antigenic relationship value (r-value) of 0.6, indicating antigenic similarity between the outbreak and vaccine virus strains. In contrast to the outbreak viruses, the antigenic relationships of the carrier viruses with the vaccine strain varied from sub-optimum (0.14) to high (0.82), and varied within and among animals and across collection times (Table 5). Three carrier-derived viruses collected from buffalo (CB2036, SB1932, and SB2017) and three viruses (CC5173, CC6646 and SC3568) collected from cattle at 3 months post outbreak had r-values of <0.3, indicating poor antigenic match with the vaccine. The remaining 18 isolates collected at 3 months post-outbreak from both convalescent and subclinical cattle and buffalo had r-values>0.3. The antigenic-relationship value tended to decrease with increasing time subsequent to the outbreak suggesting lower vaccine matching at later dates in at least in 11 isolates. However, in four animals a virus collected at a later time point had a higher r-value than a virus collected earlier in the study (Table 5).

**Table 5.**
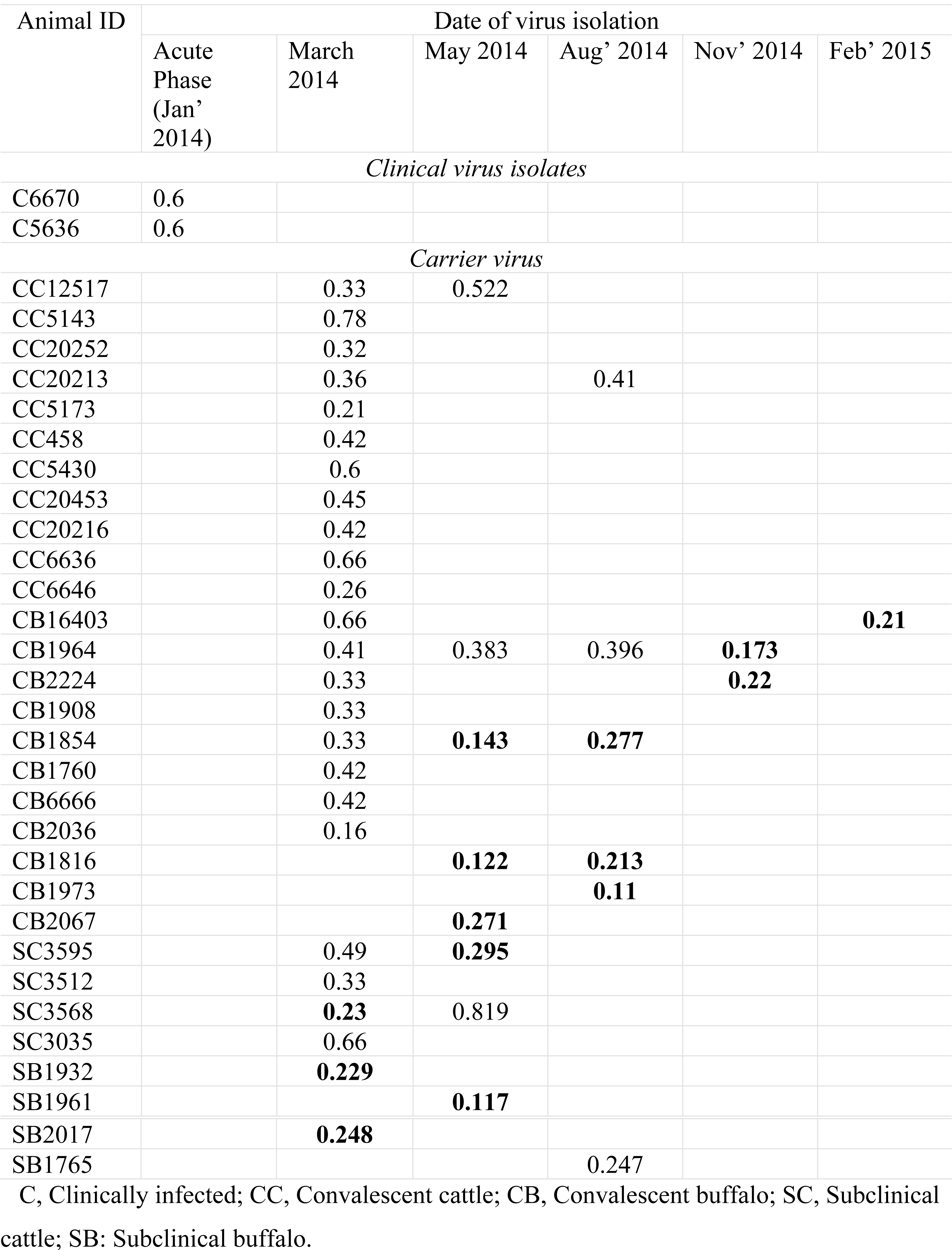
One way antigenic relationship (r-value) of the outbreak and carrier virus isolates with the currently used vaccine strain, O/IND/R2/1975, as determined by 2D-VNT assay. Values <0.3 are in bold, indicating poor vaccine protection against that isolate.

| Animal ID | Date of virus isolation |  |  |  |  |  |
| --- | --- | --- | --- | --- | --- | --- |
|  | Acute Phase (Jan' 2014) | March 2014 | May 2014 | Aug' 2014 | Nov' 2014 | Feb' 2015 |
| *Clinical virus isolates* |  |  |  |  |  |  |
| C6670 | 0.6 |  |  |  |  |  |
| C5636 | 0.6 |  |  |  |  |  |
| *Carrier virus* |  |  |  |  |  |  |
| CC12517 |  | 0.33 | 0.522 |  |  |  |
| CC5143 |  | 0.78 |  |  |  |  |
| CC20252 |  | 0.32 |  |  |  |  |
| CC20213 |  | 0.36 |  | 0.41 |  |  |
| CC5173 |  | 0.21 |  |  |  |  |
| CC458 |  | 0.42 |  |  |  |  |
| CC5430 |  | 0.6 |  |  |  |  |
| CC20453 |  | 0.45 |  |  |  |  |
| CC20216 |  | 0.42 |  |  |  |  |
| CC6636 |  | 0.66 |  |  |  |  |
| CC6646 |  | 0.26 |  |  |  |  |
| CB16403 |  | 0.66 |  |  |  | **0.21** |
| CB1964 |  | 0.41 | 0.383 | 0.396 | **0.173** |  |
| CB2224 |  | 0.33 |  |  | **0.22** |  |
| CB1908 |  | 0.33 |  |  |  |  |
| CB1854 |  | 0.33 | **0.143** | **0.277** |  |  |
| CB1760 |  | 0.42 |  |  |  |  |
| CB6666 |  | 0.42 |  |  |  |  |
| CB2036 |  | 0.16 |  |  |  |  |
| CB1816 |  |  | **0.122** | **0.213** |  |  |
| CB1973 |  |  |  | **0.11** |  |  |
| CB2067 |  |  | **0.271** |  |  |  |
| SC3595 |  | 0.49 | **0.295** |  |  |  |
| SC3512 |  | 0.33 |  |  |  |  |
| SC3568 |  | **0.23** | 0.819 |  |  |  |
| SC3035 |  | 0.66 |  |  |  |  |
| SB1932 |  | **0.229** |  |  |  |  |
| SB1961 |  |  | **0.117** |  |  |  |
| SB2017 |  | **0.248** |  |  |  |  |
| SB1765 |  |  |  | 0.247 |  |  |

C, Clinically infected; CC, Convalescent cattle; CB, Convalescent buffalo; SC, Subclinical cattle; SB: Subclinical buffalo.

## Discussion

Foot-and-mouth disease is an economic burden on endemic countries, primarily due to trade restrictions imposed by FMD-free countries in response to the risk of transmission from acutely infected animals and contaminated products. The risk associated with persistently infected FMDV carriers remains controversial, yet trade policies effectively treat carriers as infectious. In relation to the issue of infectiousness, there is also a potential risk that within-host evolution of FMDV strains during persistent infection may result in new virus variants which may subvert host immunity to cause new outbreaks. The current study characterized antigenic and genetic variation of naturally occurring FMDV O/ME-SA/Ind2001d persistent infection amongst vaccinated dairy cattle and buffalo in India. Furthermore, we demonstrated direct evidence of decreased antigenic matching of FMDVs recovered during the carrier phase under natural conditions.

In the current study, the majority of persistently infected cattle and buffalo cleared the infection by 13 months post-infection with some variation due to host species and detection of FMDV RNA vs. infectious virus. This study supports previous reports that most cattle clear persistent infections by 12 months post infection (54, 55). Interestingly, in the current study we detected a significant decrease in the proportion of persistently infected animals, based on FMDV RNA detection, between 10 and 13 months post-outbreak, which was similar to a previous report of an O/ME-SA/Ind2001d outbreak on two distinct dairy farms in India (16). However, the decrease in the proportion of persistently infected animals was reported to be more gradual for persistently infected animals under experimental conditions (15) and another field study (17). The significant decrease in the proportion of carrier animals between 10-13 months in this study may be due to differences in virus strains or host factors (vaccination status, husbandry, species) compared to previous studies. Alternatively, the decrease noted in this study may be an artefact of sample handling or laboratory artefacts.

Similar to previous reports, there was no difference in the proportion of carriers or in the duration of persistence between convalescent and subclinically infected animals (16, 55). Additionally, the proportion of animals from which FMDV RNA was recovered was similar between cattle and buffalo, however infectious FMDV was isolated from a higher proportion of animals, and for a longer duration in buffalo compared to cattle. This may reflect differences in host-virus interactions between cattle and buffalo that may enable longer survival in buffalo. Alternatively, differences in secretory antibodies (avidity or quantity) between cattle and buffalo may decrease the successful recovery of infectious virus from cattle samples. However, the small sample sizes in this study preclude definitive interpretation of this finding. Interestingly, FMDV RNA was recovered from a greater proportion of samples than virus isolates. Previous studies have shown a higher sensitivity of PCR compared to VI (14, 56, 57), and the results of the current study may reflect similar differences in sensitivity. However, other studies have reported similar sensitivities for PCR and VI (58). Overall, multiple biological and artefactual phenomena may contribute to the relative efficacies of viral detection by rRT-PCR and VI. Additional studies of naturally infected herds are needed to further characterize FMDV persistent infection under endemic conditions and differences between cattle and buffalo.

The viruses recovered in the current study aligned within the O/ME-SA/IND2001d sublineage in maximum likelihood phylogenetic analysis. Interestingly, sequences from cattle-derived isolates clustered separately from buffalo-derived isolates, suggesting the potential of host-defined, species-specific selection pressure upon FMDV evolution. An alternative interpretation is that the differential clustering of cattle and buffalo derived viruses may reflect the spatial separation of the two species in different pens within the same farm.

While classical phylogenetic analyses, such as maximum likelihood, cluster closely related virus isolates together and statistically infer ancestral relationships, the putative origin of each isolate and the genealogical relationships between the isolates cannot always be ascertained by these methods. In order to complement the conventional phylogenetic analyses, we used statistical parsimony analysis to further investigate relationships among sequences obtained in this study. Unlike phylogenetic analyses, statistical parsimony can test whether some sequences included in the analysis are ancestral to others, and in the current study all of the carrier-derived viruses originated from the outbreak virus. Similar to the phylogenetic analysis, buffalo-derived isolates clustered separately from cattle-derived isolates in the parsimony analysis. Interestingly, buffalo-derived isolates descended in a single lineage from the outbreak virus, whereas cattle-derived viruses descended from the outbreak virus in 5 separate lineages, suggesting differential selection pressures between host species. In general, viruses accumulated more SNPs over successive time points, thereby diverging from the outbreak viruses at the root node. Viruses collected from the same animal at successive time points tended to be most closely related to each other, however the later samples were not directly descended from the earlier samples.

The mechanisms driving the within-host evolution cannot be definitively determined from this study, but likely represent a combination of factors. The high divergence noted in some isolates may be due to point mutation and/or emergence of sub-consensus (minority) FMDV haplotypes from the heterogeneous populations (quasispecies) in the carrier animals, as have been described for FMDV and other picornaviruses (12, 29, 59, 60). Interestingly, three buffalo (CB1964, CB2224, CB16403) from which multiple sequences were obtained had one sequence located in a cluster distinct from the other(s), and these groupings were also supported by phylogenetic analyses. Because animals were co-habitating and sharing physical resources with imperfect biosecurity, it is possible that some apparent evolutionary changes may represent neoteric superinfection by viruses moving between animals within the herd, as has been described in buffalo in Pakistan (14). Overall, the statistical parsimony analysis suggested that carrier isolates followed distinct routes of evolution in different FMDV-persistently infected individuals and species. This may suggest that species-defined and/or individual animal-level selective pressures may have determined the evolutionary paths.

The current study attempted to test the hypothesis that during natural persistent FMDV infection, substitutions would occur at specific regions of the capsid coding region, reflecting viral mechanisms of immune escape and maintenance of persistence. Two codons under positive selection in this study were in VP1 (138 & 148) and are located on the VP1 βG-βH loop, while one codon was in VP2 (VP2-78) and is located close to antigenic site-2. Although one site with a synonymous substitution was present in 5 out of 8 carrier animals, a consistent pattern of amino acid changes amongst all carrier isolates was not detected. Previous studies have addressed this subject with variable results. One study identified consistent change in the B–C loop of VP2 during persistence in cattle (19), whereas another study identified a pattern of Q-172-R substitution in VP1 (20). Other studies have concluded that a consistent pattern of substitution does not occur during the carrier state (25). These results suggest that viral determinants may have some role, but are not likely to be solely responsible for determining persistent FMD infection or for FMDV evolution within carrier animals. The selection pressure acting on the viral genome varies among individual persistently infected animals, suggesting that host factors are similarly important to the viral determinants, as has been suggested by some authors (61, 62).

Few studies have investigated how antigenicity changes during persistent infection. In the current study, the antigenic-relationship (r-value) with the vaccine strain decreased over time, often below the threshold of protection. This accentuates the finding of antigenic divergence reported herein as it represents one of very few times where direct evidence is found of decreasing antigenic matching during FMDV persistence. This contributes to understanding the underlying evolutionary mechanisms for the emergence of new strains during viral persistence. However, r-value is not a perfect indicator of cross-protection as there are many examples in which vaccine protection *in vivo* did not correlate with vaccine matching performed *in vitro* (63). Despite the limitations of r-determinations, the decreasing trend in the current study is concerning because of the potential for these viruses to cause new outbreaks, even in vaccinated animals, if transmitted.

The evolutionary rate estimated in this study was similar to the rate reported for other serotype O carrier viruses (2.6 × 10^-2^ s/s/y;(25)).In contrast, the rate reported in this study was an order of magnitude faster than the rate reported for O/ME-SA/Ind2001 outbreak isolates collected in India between 2000 – 2013 (6.338 × 10^-3^s/s/y;(41)). Similarly, previous studies have shown the rate of evolution of serotype C carrier viruses was an order of magnitude faster than the rate reported for serotype C outbreak viruses collected over a period of six decades (59, 64). The FMDV genome appears to be under higher selection pressure during persistent infection, resulting in the generation of genetic and antigenic variants. Yet, across animals, the ω value varied from 0.076 to 0.357, indicating different extent of selection pressures acting on viral genomes in different individuals. Interestingly, there did not appear to be a relationship between selection pressure and duration of persistent infection in the small number of animals in the current study.

## Conclusions

The current study contributes to elucidation of within-host evolution of FMDV in the transition from acute to carrier phases and over the course of viral persistence. Whether an animal had clinical FMD during the acute phase of infection did not affect within-host virus evolution or the dynamics of persistent infection. Overall, the genetic variation of carrier viruses presented in this study is consistent with the complexity and dynamics of *in vivo* FMDV quasispecies, and this study suggests different host species may exert differential influences which contribute to within-host viral evolution. However, variation across viruses from distinct animals precluded identification of specific mutations that define the carrier state. The antigenic-relationship of virus isolates to the vaccine strain tended to decrease during persistent infection, indicating the potential of emergence of divergent antigenic strains from carrier animals. However, the probability of transmission of these viral variants from persistently infected animals to susceptible animals, and their fitness to cause clinical FMD require further investigation. To our knowledge, this is the first report on both genetic and antigenic variation of FMDV during virus persistence in infected cattle and domestic Asian buffaloes under natural conditions.

## Acknowledgements

We are thankful to the supporting staff of the dairy farm of their cooperation during the sample collection. Technical assistance of Mr N.S. Singh and Mr B. Das are highly acknowledged. Miranda Bertram and Barbara Brito were recipients of Plum Island Animal Disease Center Research Participation fellowships, administered by the Oak Ridge Institute for Science and Education through an inter-agency agreement with the US Department of Energy. This work was funded in part by the U.S. Department of Agriculture, Agricultural Research Service-CRIS project 1940-32000-061-00D and by the Biosecurity Engagement Program of the US Department of State. The funders had no role in study design, data collection and analysis, or the decision to publish the work.

